## Supplemental Materials for "Pulsatile electrical stimulation creates predictable, correctable disruptions in neural firing"

### SUPPLEMENTARY INFORMATION

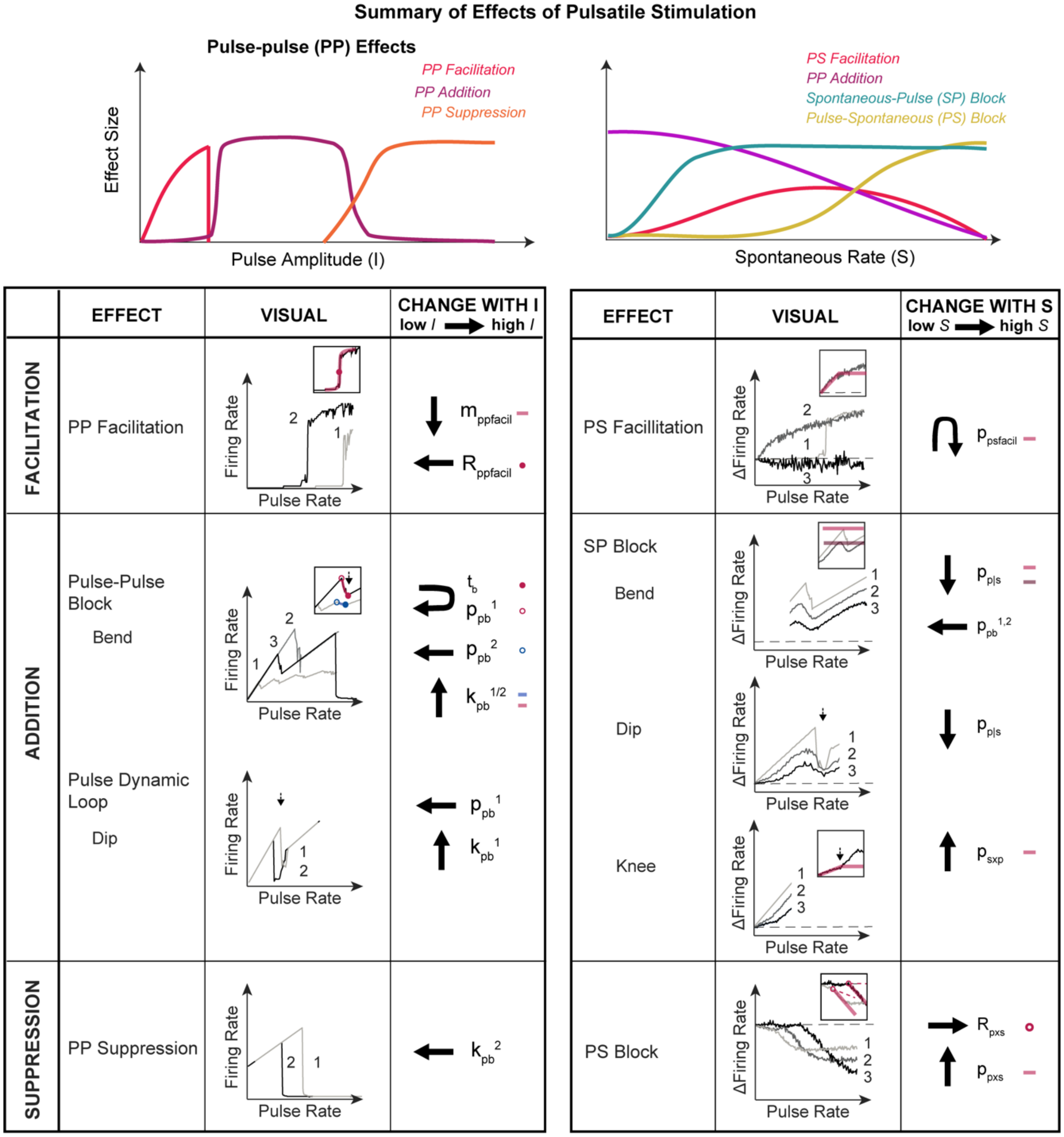


***Supplemental Figure 1. Summary of Effects of Pulsatile Stimulation.*** *(top) Overview of Pulse Effect and how they change with Increasing Pulse Amplitude (I) (left) and Increasing Spontaneous Activity (S) (right). The traces show the rough change in each effect with I and S. See Supplemental Fig. 2 for details of how each parameter is affected by I and S. (bottom) Three categories of effects are observed: facilitation, addition, and suppression. The main pulse-pulse effects are shown on the left and the main pulse-spontaneous effects are shown on the right. In each table, the name of each effect is provided as described in the text. The center panel highlights the change in shape of the pulse rate-firing rate relationship(PFR) induced by that effect. Darkening of the PFR and increasing number next to PFR indicates higher I/S in the left/right table. The right column shows how the parameters in the predictive equation change with increasing I/S. Arrows indicate how the feature of the PFR moves with that parameter as I/S increases. The insets highlight the specific feature of the PFR being controlled by each parameter. The rule contributing to that feature is shown in red in all cases except for the case of the parameter controlling bend 1 versus bend 2 where parameters for*

*1 are in red and for 2 are in blue. In the PS block case, $p_{sxp}$ is still in effect at low S (marked 1), so $p_{pxs}$ controls the change in slope compared to the previous negative slope due to $p_{sxp}$. That reference slope is indicated in the inset with a dashed pink line.*

***Supplemental Table 1. Variables used in Equations for Modeling Pulsatile Interactions.*** Variable explanations and bounds are provided. Details of how each parameter is affected by stimulation conditions and in turn affects the PFR can be visualized in Supplemental Figures 1 & 2.

| Variable | Bounds | Meaning |
|---|---|---|
| **F** | -- | Induced firing rate (in the presence of spontaneous firing, pulses, etc.) |
| **R** | -- | Pulse rate |
| **I** | -- | Pulse amplitude |
| $\mathbf{I_{Na}}$, $\mathbf{I_{KH}}$, $\mathbf{I_{KL}}$, **etc.** | -- | Current for each of the channels driving change in axonal state |
| **S** | -- | Spontaneous firing rate of neuron under no stimulation |
| $\mathbf{I_{pred}}$/ $\mathbf{F_{pred}}$ | -- | Predicted pulse amplitude estimated by minimizing the error between data and model prediction for induced firing rate at a given current amplitude. $F_{pred}$ – corresponding firing rate for a prediction |
| $\mathbf{F_{pp}}$ | -- | Contribution to induced firing rate of spontaneous activity-induced effects |
| $\mathbf{F_{ss}}$ | -- | Contribution to induced firing rate of pulse-induced effects |
| $\boldsymbol{\psi}_1$ | -- | Condition on first step with partial elimination under extreme pulse amplitudes. (Pulse dynamic loop Rule where there is exponential decay between $t_{pb}$ and $t_b$.) |
| $\boldsymbol{\psi}_2$ | -- | Condition on second step with partial elimination under extreme pulse amplitudes. (Suppression of Future Pulses where there is exponential decay toward zero action potentials being produced.). |
| ***n*** | [1,2] | The number of the step in the pulse-rate firing rate relationship being considered. The same parameters as for *n*=2 were used for *n* >2. |
| $\mathbf{t_b}$ | [1:600] | The time after a pulse in which pulses produce action potentials with probability less than 0. |
| $\mathbf{p_{pb}^n}$ | (0:1) | The portion of $t_b$ that the partial elimination effect lasts, where pulses produce action potentials with probability less than 1. One per *n.* |
| $\boldsymbol{\kappa}_{\mathbf{pb}}^{\mathbf{n}}$ | (0,600) | The scaling of the partial block window. One per *n.* |
| $\mathbf{p_{sxp}}$ | [0:1] | Probability of spontaneous EPSCs having a significant interaction with pulses that blocks a pulse-induced AP |
| $\mathbf{p_{pxs}}$ | [0:1] | Probability of a pulse having a significant interaction with spontaneous EPSCs that blocks a spontaneous AP |
| $\mathbf{R_{pxs}}$ | [0:600] | The pulse rate at which pulses begin to block spontaneous APs |
| $\mathbf{p_{p\|s}}$ | [0:1] | Probability of pulses forming an action potential given the state of the axon and existing EPSCs |
| $\mathbf{p_{psfacil}}$ | [0:1] | Probability of pulses being facilitated to make APs when they alone are incapable due to the presence of EPSCs |
| $\mathbf{m_{ppfacil}}$ | (0:1] | The slope of the facilitation starting when pulse-pulse facilitation occurs (or there is no spontaneous activity). |
| $\mathbf{R_{ppfacil}}$ | [0:600] | The pulse rate at which the facillitation is centered when pulse-pulse facilitation occurs. This is where a sudden jump to pulses producing APs occurs. |

***Supplemental Table 2. RMS across all amplitudes for each Simulated Spontaneous Firing Rate:*** The average rms across all fits was 5.77 ± 1.19 sps.

| **Spontaneous Firing Rate (*sps*)** | **RMSE Cross Amplitudes (sps)** | **T-test Results Fitted v. Unfitted Amplitudes** |
|---|---|---|
| 0 | 1.93 ± 0.67 | t(42)= 0.19, p=0.86 |
| 5 | 4.19 ± 1.22 | t(42)= -0.56, p=0.58 |
| 13 | 4.64 ± 0.90 | t(42)= -1.14, p=0.26 |
| 28 | 4.99 ± 0.62 | t(42)=-1.48, p=0.15 |
| 51 | 5.53 ± 0.44 | t(42)=-0.23, p=0.82 |
| 80 | 7.14 ± 0.40 | t(42)=-1.59, p=0.12 |
| 131 | 11.98 ±0.69 | t(42)=-1.18, p=0.24 |

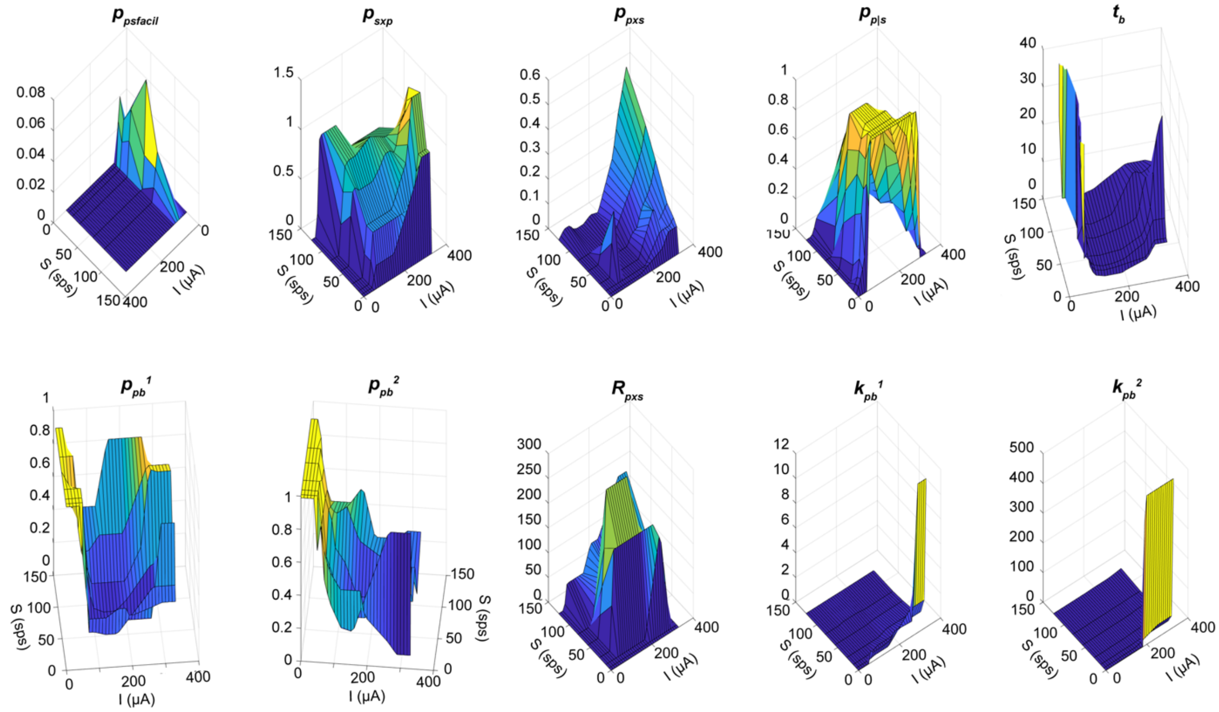


***Supplemental Figure 2. Depedence of Equation Parameters on Pulse Amplitude and Spontaneous Rate.*** *A plot of the dependence of each variable used in the pulsatile stimulation prediction equations ($p_{psfacil}$, $p_{sxp}$, $p_{pxs}$, $p_{p|s}$, $t_b$, $t_{pb}^1$, $t_{pb}^2$, $R_{pxs}$, $k_{pb}^1$, $k_{pb}^2$) on pulse amplitude (I) and spontaneous rate (S).*

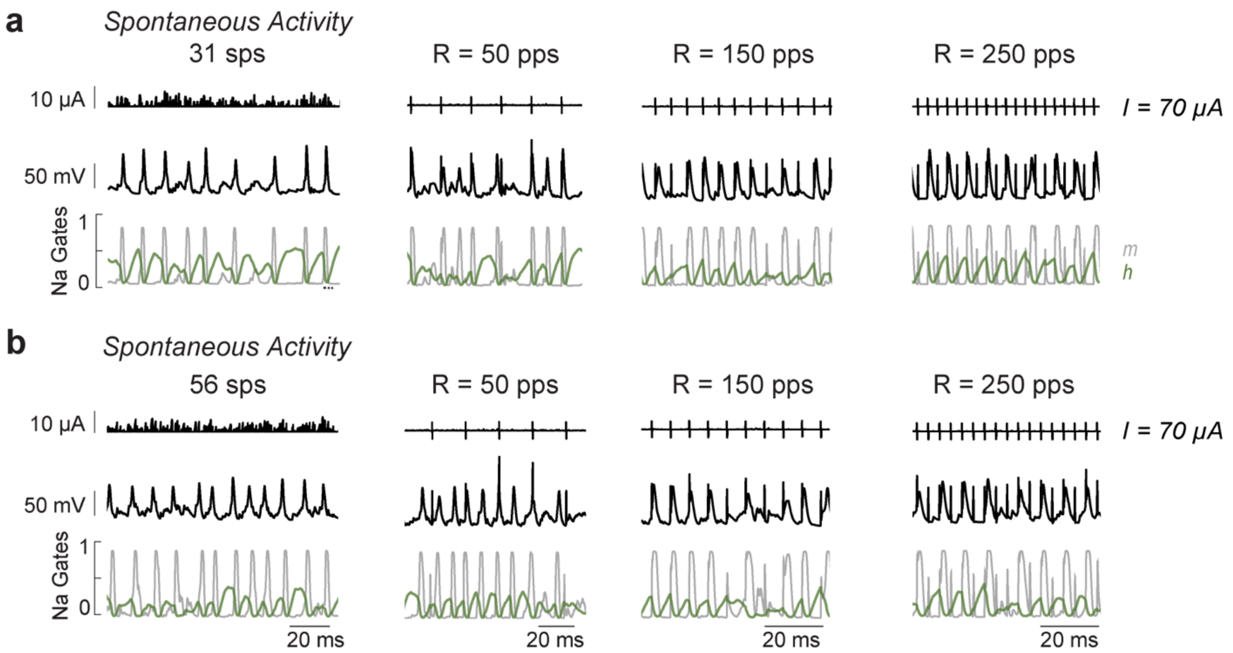


***Supplemental Figure 3. Pulse Effects with Spontaneous Activity.*** *Channel dynamics during spontaneous activity and pulse-spontaneous interactions. External current input (top), voltage trace (middle), sodium channel dynamics (bottom) with m-gate(grey) and h-gate(green). On the left are responses with no pulsatile stimulation, and the plots to the right are the same simulated afferent responding to R= 50 pps, 150 pps, and 250 pps. a) Responses for afferent with a spontaneous rate of 31 sps. b) Responses for afferent with a spontaneous rate of 56 sps.*

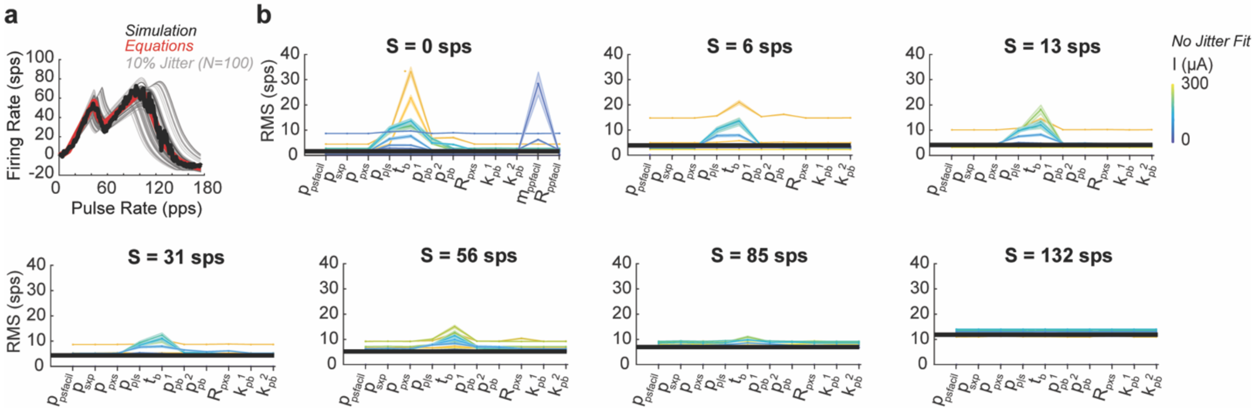


***Supplemental Figure 4. Prediction Equation Parameter Sensitivity Analysis.*** *a) Example of 100 iterations of 10% jitter of $t_b$. The simulation (black) is compared to the optimal fit (red), and jitters(grey). b) Plot per spontaneous rate (S) of how a 10% jitter of each parameter affects rms between equation fit and simulation at each current amplitude between 0 and 300 (colored as in graph). Pre-jitter rms value shown in black.*

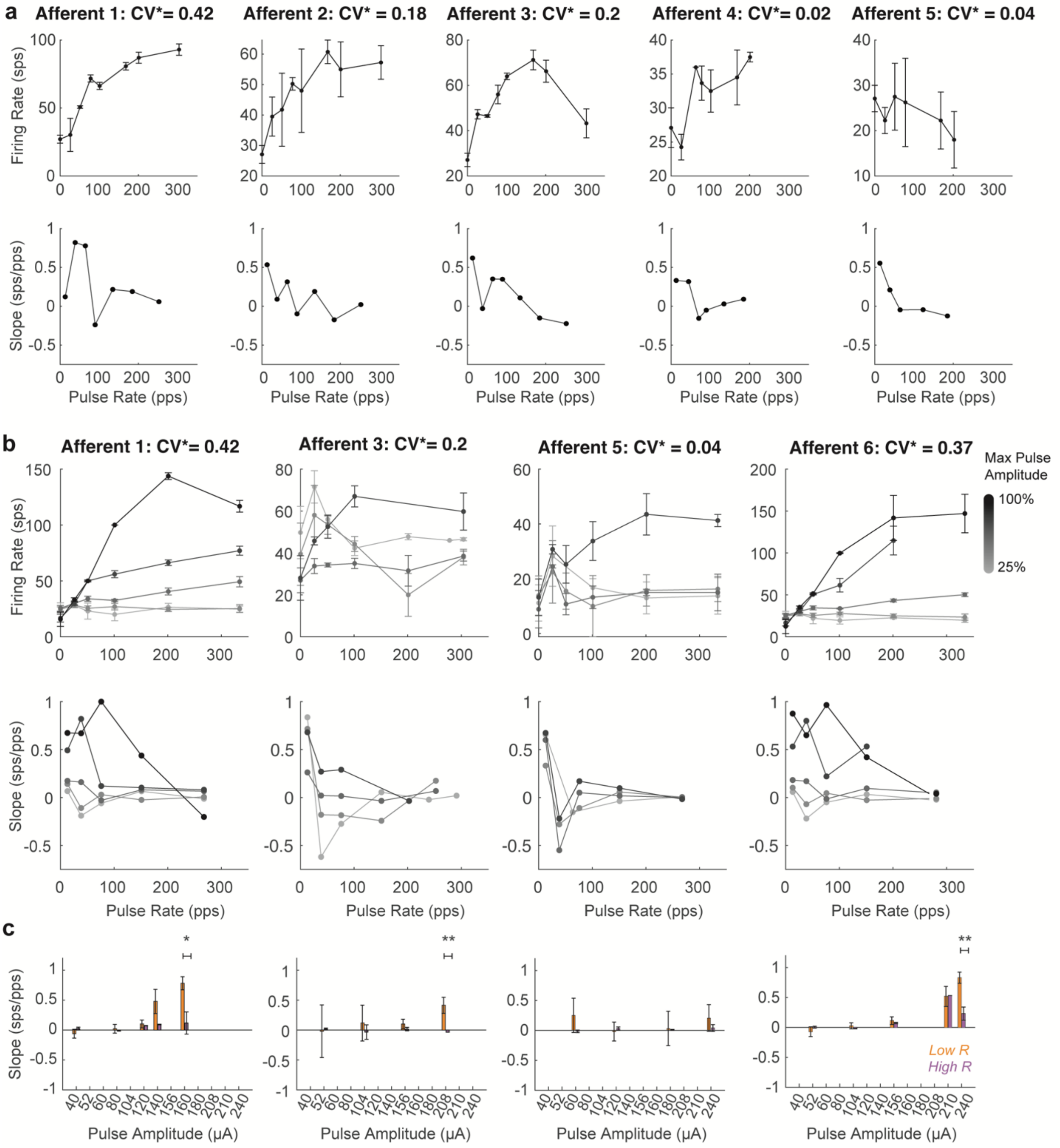


***Supplemental Figure 5. Experimental Afferent Responses to Pulsatile Stimulation.*** *a) Individual recordings from five afferents at 80% of peak safe current amplitude, where safe current amplitude per subject was level of facial twitch. b) Recordings from four afferents three of which overlap A) as current was stepped up from 25% to 100% of the safe range of amplitudes per animal where the highest amplitude was 250 µA. Below is the slope between each PFR sampling step. c) Low R vs. high R slope prevalence with SEM per afferent with amplitude. Significant differences observed at different amplitudes (unpaired t-test:*, p<0.1;** p<0.05). Statistical power was over 0.85 in all cases with*.*

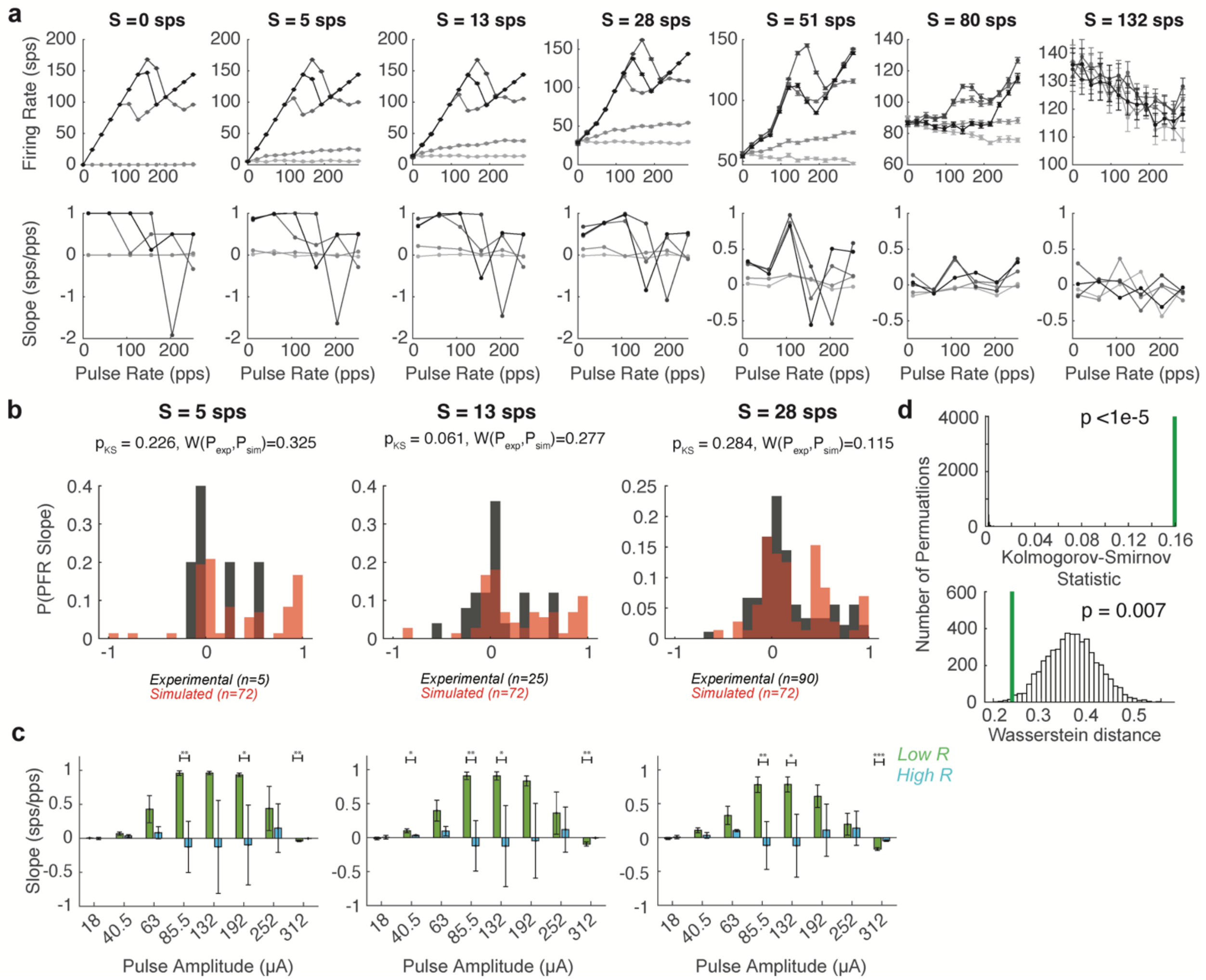


***Supplemental Figure 6. Simulated Responses Compared to Experimental Results**. a) Simulation of PFR at different spontaneous rates with sparse sampling of the PFR every 30 pps. (below) Slopes with increased current show in grayscale with highest pulse amplitude in black. Only pulse rates from 0-300 sps were used for comparability to recorded data. b) PDFs of slopes at each spontaneous rate across all experiments (black) compared to simulated slopes for afferents within the same spontaneous rate category (red), where categories are centered at simulated Ss.* For some groups, this represents only one afferent. *Kolmogorov-Smirnov statistic ($p_{KS}$) and Wasserstein distance $W(P_{exp},P_{sim})$ reported above each plot. From left to right unpaired t-test results were insignificant with (t(75)=0.53, p=0.60; t(95)=1.32, p=0.19; t(160)=0.24, p=0.81). c) Low R versus high R slope per sampled pulse amplitude for the same spontaneous rate group (unpaired t-test:*, p<0.1;** p<0.05;***, p<0.01). d) Comparison of Kolmogorov-Smirnov statistic and Wasserstein distance overall to values for slopes of 5000 permutations of pulse rate-firing rate combinations. Reported p-value for each is rank of original experimental count compared to 5000 permutations.*

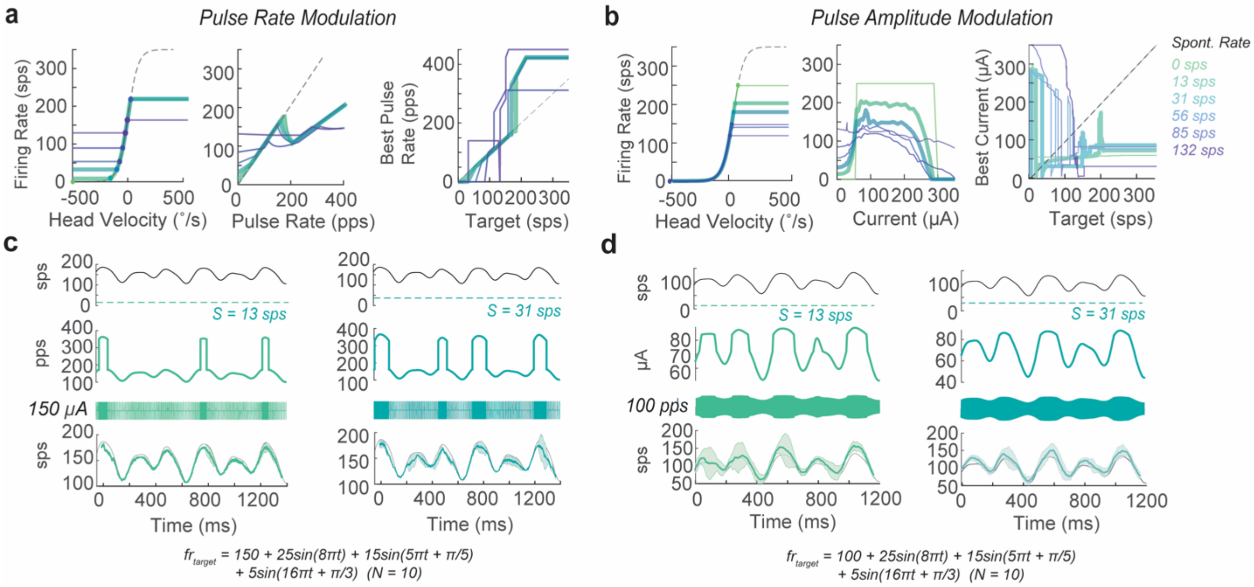


***Supplemental Figure 7. Pulsatile Stimulation Rules during Pulse Modulation.*** *Optimal pulse parameters per simulated spontaneous rate, with same colors as in Fig. 3-4. (left) Maximum restoration of the firing rate encoding compared to head velocity range. (middle) maximal PFR for that condition. (right) target firing rate to best pulse parameter mapping, considering minimizing the pulse parameter. a) best parameters for PRM, I = 250 μA. b) Best parameters for PAM, R=100 pps. More complex target firing patterns that were a mixture of sinusoids in the physiological range were tried. Equations below c-d. Example optimal parameters for the same complex target firing pattern for afferents with different spontaneous rate firings rates for a c) PRM and d) PAM strategy, shown in the same way as in Fig. 5c.*

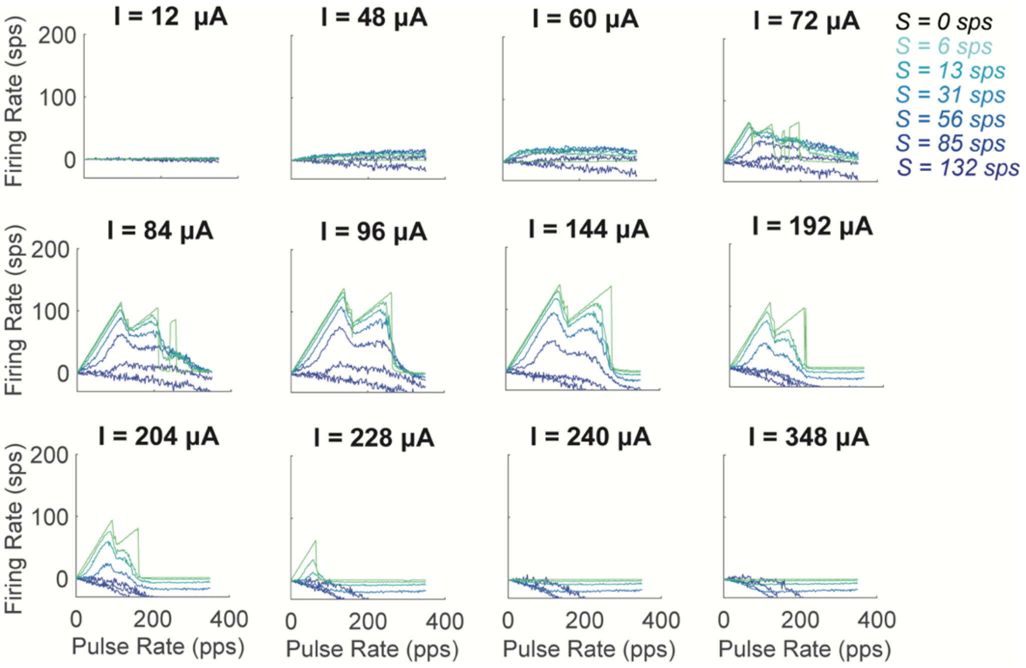


***Supplemental Figure 8. Effect of Conductance on Response with Increase Amplitude.*** *Mapping of PFRs for same range of baseline spontaneous rates 0 to 131 sps. We note a quick transition from facilitation to pulse-spontaneous blocking at lower current amplitudes.*

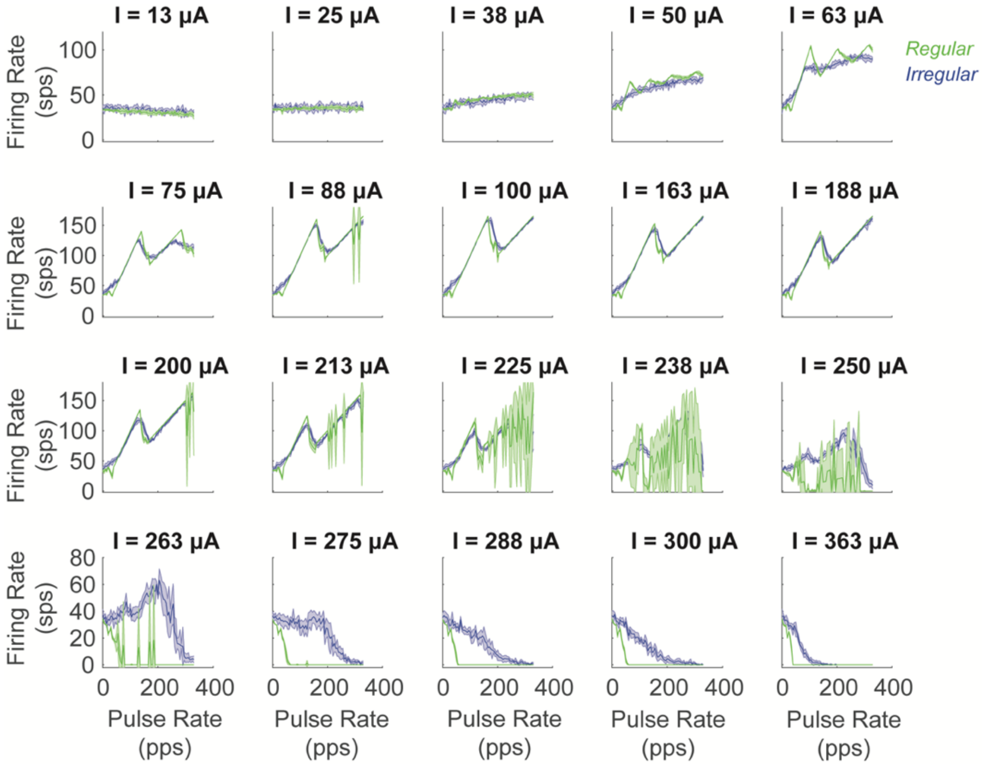


***Supplemental Figure 9. Effect of Regularity on Pulsatile Stimulation****. Mapping of PFR for I from 0 – 363 μA for regular neuron (green, CV = 0.09) and irregular neuron (blue, CV = 0.57) with similar firing rates (32-36 sps). The response at the same current amplitude is plotted for 5 repetitions, showing differences in the variance at different amplitudes and the magnitude of the reduced length of the PE zone for regular neurons. The start of blocking is observed to start at lower I for regular neurons. Additionally, when R < S the response dips, indicating a stronger spontaneous-pulse block.*

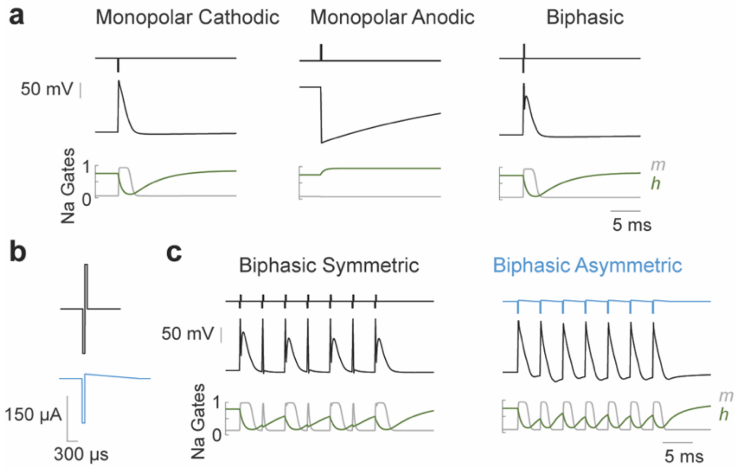


***Supplemental Fig 10. Shaping Pulses to Overcome Blocking Effects****. Simulations are shown for each of the following: a) A monopolar cathodic pulse, monopolar anodic pulse, and biphasic pulse made from the combination of the two and the voltage change (black) and channel effect, specifically on h(green) and m(grey) that results. The cathodic and biphasic pulse produces highly similar channel dynamics and voltage changes resembling an AP. b) (top) A standard symmetric waveform compared to (blue) an asymmetric waveform with an anodic phase designed to overcome blocking effects by creating a small afterhyperpolarization. Both are the same amplitude and charge-balanced and were delivered at the same rate. c) The results of each waveform on channel dynamics and voltage. Biphasic asymmetric pulses restore a one-to-one PFR instead of causing pulse-pulse blocking.*
